# Anesthetic agents affect urodynamic parameters and anesthetic depth at doses necessary to facilitate preclinical testing in felines

**DOI:** 10.1101/868398

**Authors:** Jiajie Jessica Xu, Zuha Yousuf, Zhonghua Ouyang, Eric Kennedy, Patrick A. Lester, Tara Martin, Tim M. Bruns

**Affiliations:** Unit for Laboratory Animal Medicine, University of Michigan, Ann Arbor, MI; Biointerfaces Institute, University of Michigan, Ann Arbor, MI; Biomedical Engineering Department, University of Michigan, Ann Arbor, MI

**Author notes:** **Corresponding Author:** Tim M. Bruns, Department of Biomedical Engineering, University of Michigan, 2800 Plymouth Road, NCRC B10-A169, Ann Arbor, MI, 48109, USA.

**Keywords:** anesthesia, bladder, cats, urodynamics

## Abstract

Urodynamic studies, used to understand bladder function, diagnose bladder disease, and develop treatments for dysfunctions, are ideally performed with awake subjects. However, in animal models, especially cats (a common model of spinal cord injury and associated bladder pathology), anesthesia is often required for these procedures and can be a research confounder. This study compared the effects of select agents (dexmedetomidine, alfaxalone, propofol, isoflurane, and α-chloralose) on urodynamic (Δpressure, bladder capacity, bladder compliance, non-voiding contractions, bladder pressure slopes) and anesthetic (change in heart rate [ΔHR], average heart rate [HR], reflexes, induction/recovery times) parameters in repeated cystometrograms across five adult male cats. Δpressure was greatest with propofol, bladder capacity was highest with α-chloralose, non-voiding contractions were greatest with α-chloralose. Propofol and dexmedetomidine had the highest bladder pressure slopes during the initial and final portions of the cystometrograms respectively. Cats progressed to a deeper plane of anesthesia (lower HR, smaller ΔHR, decreased reflexes) under dexmedetomidine, compared to propofol and alfaxalone. Time to induction was shortest with propofol, and time to recovery was shortest with dexmedetomidine. These agent-specific differences in urodynamic and anesthetic parameters in cats will facilitate appropriate study-specific anesthetic choices.

## Introduction

Urodynamic studies such as cystometrograms (CMGs) are used to study bladder function, diagnose bladder disease, and even develop treatments for bladder dysfunctions. Ideally, the subject should be awake and unanaesthetized during these studies ^1^. However anesthesia, which can confound urodynamic outcomes ^2-4^, is often required in many animal models of urodynamic studies, particularly when treatments for bladder dysfunctions are being tested in experimental paradigms that require control over the animal’s state.

Cats are a common animal model for spinal cord injury and associated bladder pathology. Despite their relatively small body size, their spinal cord is of similar length and anatomy to the human spinal cord ^5-7^. Urodynamic studies in awake cats are challenging. Placement of indwelling bladder catheters for awake studies is invasive, and can lead to complications such as cystitis. Temporary catheter placement for urodynamic trials necessitates the administration of anesthetic agents ^8^ that interfere with autonomic reflexes to varying degrees ^9,10^. Agents minimally inhibitory towards autonomic reflexes such as α-chloralose or urethane, are not recommended for survival procedures due to concerns of prolonged recovery or carcinogenicity^11^.

While current literature has shown differences in CMG parameters such bladder capacity, peak pressure, and non-voiding contractions (NVCs) based on different anesthetic agents, studies are limited especially in cats. To our knowledge, no studies have characterized the following agents used in clinical veterinary medicine for cats on CMG parameters: 1) alfaxalone, 2) dexmedetomidine, and 3) propofol (without the use of another anesthetic).

Dexmedetomidine is commonly used as a sedative-anesthetic in veterinary medicine (and to facilitate CMG testing in our laboratory ^12^), due to its ease of administration (intramuscular [IM], and rapid recovery through the reversal agent atipamezole). However, in a rat study, dexmedetomidine was inhibitory to the volume evoked micturition reflex and peak pressure compared to urethane and awake trials ^13^. Propofol, an anesthetic agent used to facilitate CMG testing in clinical veterinary medicine ^14,15^, has not been compared directly to dexmedetomidine in cat trials, but is suspected to be less inhibitory than dexmedetomidine due to its mechanism of action (propofol has no analgesic properties, while dexmedetomidine does) and ability to carefully titrate anesthetic depth through intravenous [IV] infusion. Unfortunately, propofol can only be given IV, so it often requires another agent to secure IV access. Alfaxalone, recently approved in the United States for use in cats, has a similar mechanism of action and clinical application as propofol, but has not been used in cats in the context of urodynamic studies. Unlike propofol, alfaxalone is formulated for IM dosing and does not require an additional agent^16^.

In this study we anesthetized male cats with alfaxalone, dexmedetomidine, and propofol at the lowest anesthetic doses practical to facilitate CMG testing prior to terminal procedures that used isoflurane and α-chloralose. We assessed the effect of anesthetic agent on CMG and anesthetic parameters, hypothesizing that some anesthetic agents would be more inhibitory towards bladder function than others. Vascular access ports (VAPs) were placed prior to trials, to ensure propofol could be studied independently without the need for another agent.

## Results

Five cats were anesthetized at least 3 times with each agent (Supplementary Table S1). At least 2 CMG trials were conducted per session. We only collected terminal (isoflurane and α-chloralose) trials from four of five cats due to one unexpected death.

### Cystometrogram Parameters

Bladder capacity differed significantly between agents (Fig. 1a, Supplementary Table S2, Supplementary Table S3). Bladder capacity was greatest with α-chloralose (59.6 ± 9.2 ml), and significantly increased compared to alfaxalone (46.7 ± 8.6 ml, p = 0.02), dexmedetomidine (42.9 ± 8.6 ml, p < 0.001), and propofol (44.2 ± 8.6 ml, p = 0.003). Bladder capacity under isoflurane (54.5 ± 9.1 ml) was significantly increased compared to dexmedetomidine (p = 0.03). Additionally, Δpressure also differed significantly between agents (Fig. 1b, Supplementary Table S2, Supplementary Table S3. Trials under propofol had the greatest Δpressure (116.5 ± 9.6 cm H_2_0), and were significantly increased compared to isoflurane (84.2 ± 13.1 cm H_2_0, p = 0.04) (Fig. 1b). Compliance did not differ significantly between agents (p > 0.05 for all pairwise comparisons, Supplementary Table S2, Supplementary Table S3).

**Figure 1:**
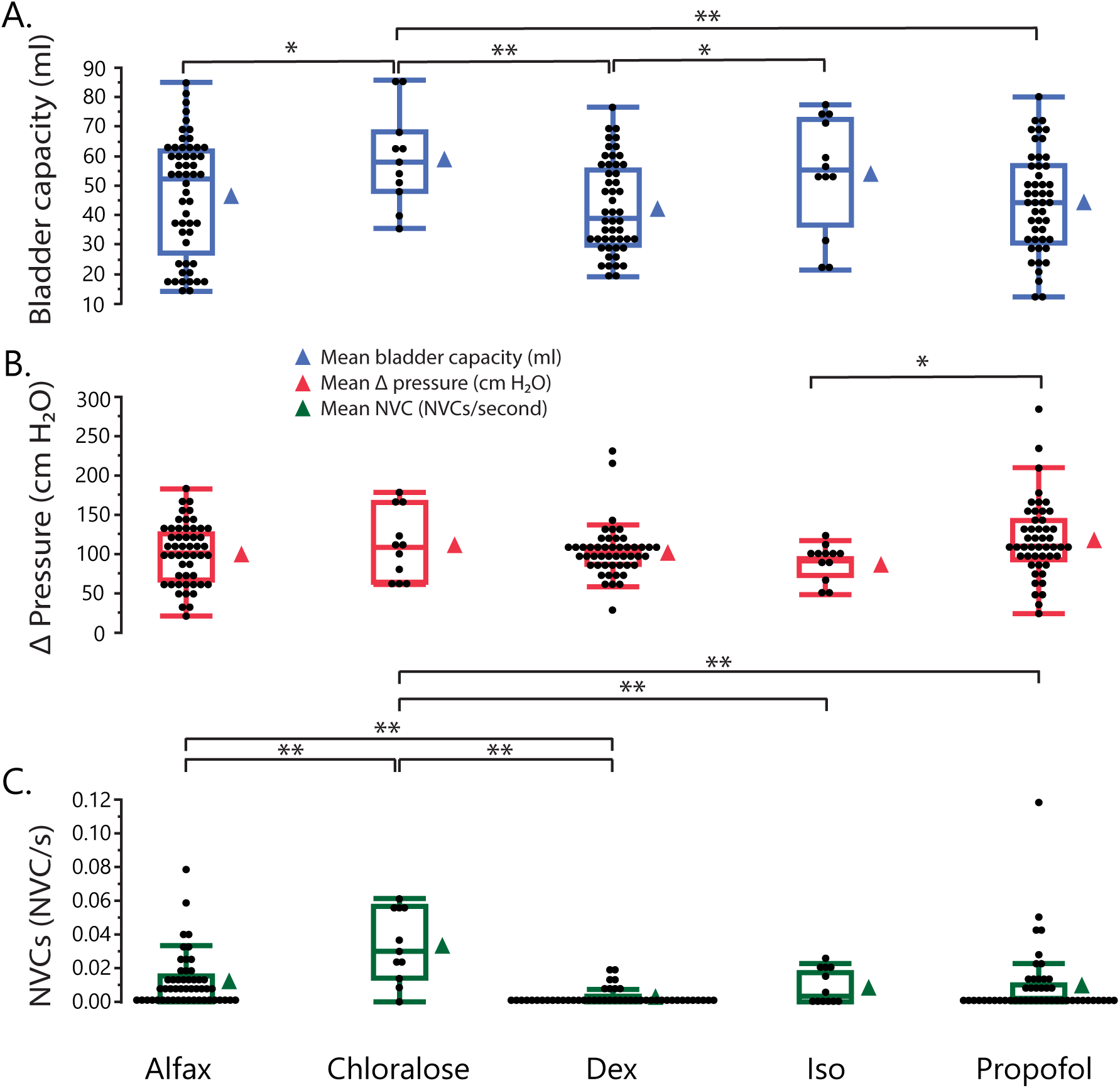
Boxplots of (A) bladder capacity, (B) Δpressure, and (C) NVC rate by anesthetic agent. Points represent individual samples. Adjacent triangle icons represent means of individual samples. Boxes represent 1^st^ to 3^rd^ quartile range (interquartile range [IQR]), with the line in the middle of each box representing the median. Tips of whiskers extending below and above box represent 1^st^ quartile – 1.5*IQR and 3^rd^ quartile + 1.5*IQR respectively. Significance: * p < 0.05, ** p < 0.01.

Non-voiding contraction rate was highest with α-chloralose (0.033 ± 0.004 NVCs/s) and significantly higher than all other agents (p < 0.01 for all comparisons; Fig. 1c, Supplementary Table S2, SupplementaryTable S3). Alfaxalone (0.012 ± 0.003 NVCs/s) had the second highest rate of NVCs, and was significantly greater than dexmedetomidine which had the lowest rate of NVCs (0.002 ± 0.003 NVCs/s, p = 0.008). The amplitude of non-voiding contractions was not different between agents (p > 0.05).

The trends of the CMG bladder pressure responses changed with agent, with different agents having variable periods of passive filling compared to active contraction over the trial. Notably, propofol CMG responses were characterized by an early and gradual rise (longer period of active contraction), compared to dexmedetomidine responses which were characterized by a long period of low pressure (passive contraction) and a rapid rise immediately before voiding (Fig. 2a, Supplementary Table S2). The bladder pressure trends for other agents fell between the propofol and dexmedetomidine extremes. This was further described through quantification of slopes at select time points (Fig. 2b). During slope 1 (start to 100 seconds before void), propofol (0.06 ± 0.01 cm H_2_0/s) was significantly increased compared to alfaxalone (0.03 ± 0.01 cm H_2_0/s, p < 0.0001) and dexmedetomidine (0.02 ± 0.01 cm H_2_0/s, p < 0.001). During slope 2 (100s before void to 50s before void), α-chloralose had the highest slope (0.51 ± 0.08 cm H_2_0/s), significantly higher than dexmedetomidine (0.17 ± 0.05 cm H_2_0/s, p = 0.001) and alfaxalone (0.27 ± 0.05 cm H_2_0/s, p = 0.03). During slope 3 (50s before void to void event) dexmedetomidine (1.45 ± 0.11 cm H_2_0/s) was significantly greater than all other agents: alfaxalone (1.10 ± 0.11 cm H_2_0/s p = 0.009), α –chloralose (0.63 ± 0.18 cm H_2_0/s, p < 0.001), isoflurane (0.57 ± 0.17 cm H_2_0/s, p < 0.001), and propofol (0.65 ± 0.11 cm H_2_0/s, p < 0.001).

**Figure 2:**
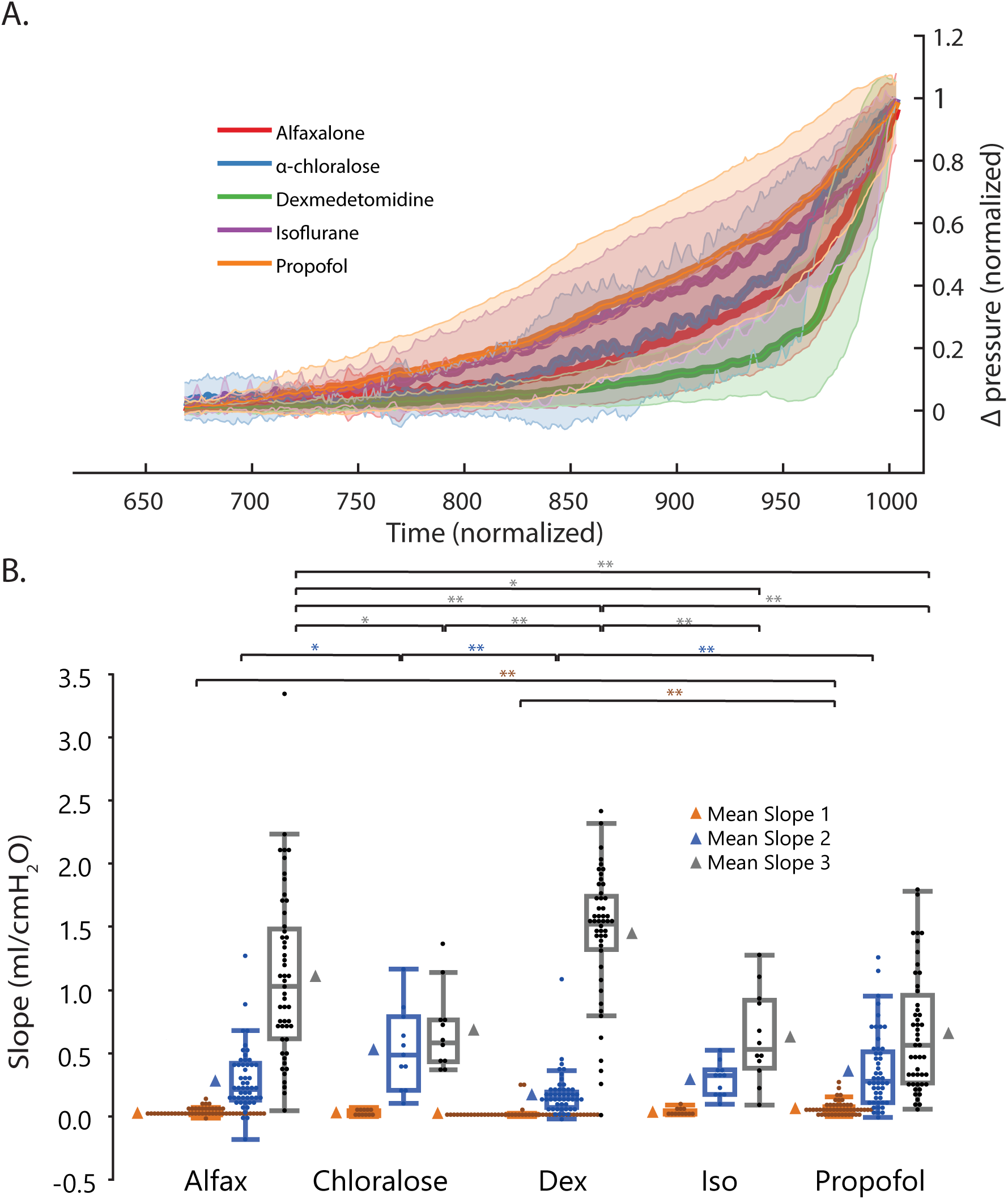
A. Normalized CMG bladder pressure responses averaged by agent. Solid lines represent means, shading around each mean represents standard deviations. B. Box plots of CMG slopes. Data represented as in Fig. 1. Significance: * p < 0.05, ** p < 0.01.

### Anesthetic Parameters

Mean HR and mean ΔHR both varied between agents (Fig. 3a, Fig. 3b, Supplementary Table S4, Supplementary Table S5). Mean HR, compared across all five agents, was highest under alfaxalone (215.16 ± 5.69 bmp), and significantly greater than all other agents: α-chloralose (163.29 ± 9.49 bmp, p < 0.001), dexmedetomidine (110.66 ± 5.75, p < 0.001), isoflurane (130.73 ± 8.18 bmp, p < 0.001), and propofol (176.72 ± 5.81 bmp, p < 0.001). All other agents fell in between the two extremes. Mean change in heart rate (ΔHR), compared across only non-terminal agents, was lowest with dexmedetomidine (13.79 ± 6.80 bmp), and significantly different from propofol (37.50 ± 7.05 bmp, p = 0.0082) and alfaxalone (45.36 ± 6.65 bmp, p < 0.001). Mean ΔHR did not differ significantly between propofol and alfaxalone (p = 0.76). Two trials (both propofol) were excluded from the averaged continuous HR plot (Fig. 3a) due to suspect artifacts.

**Figure 3:**
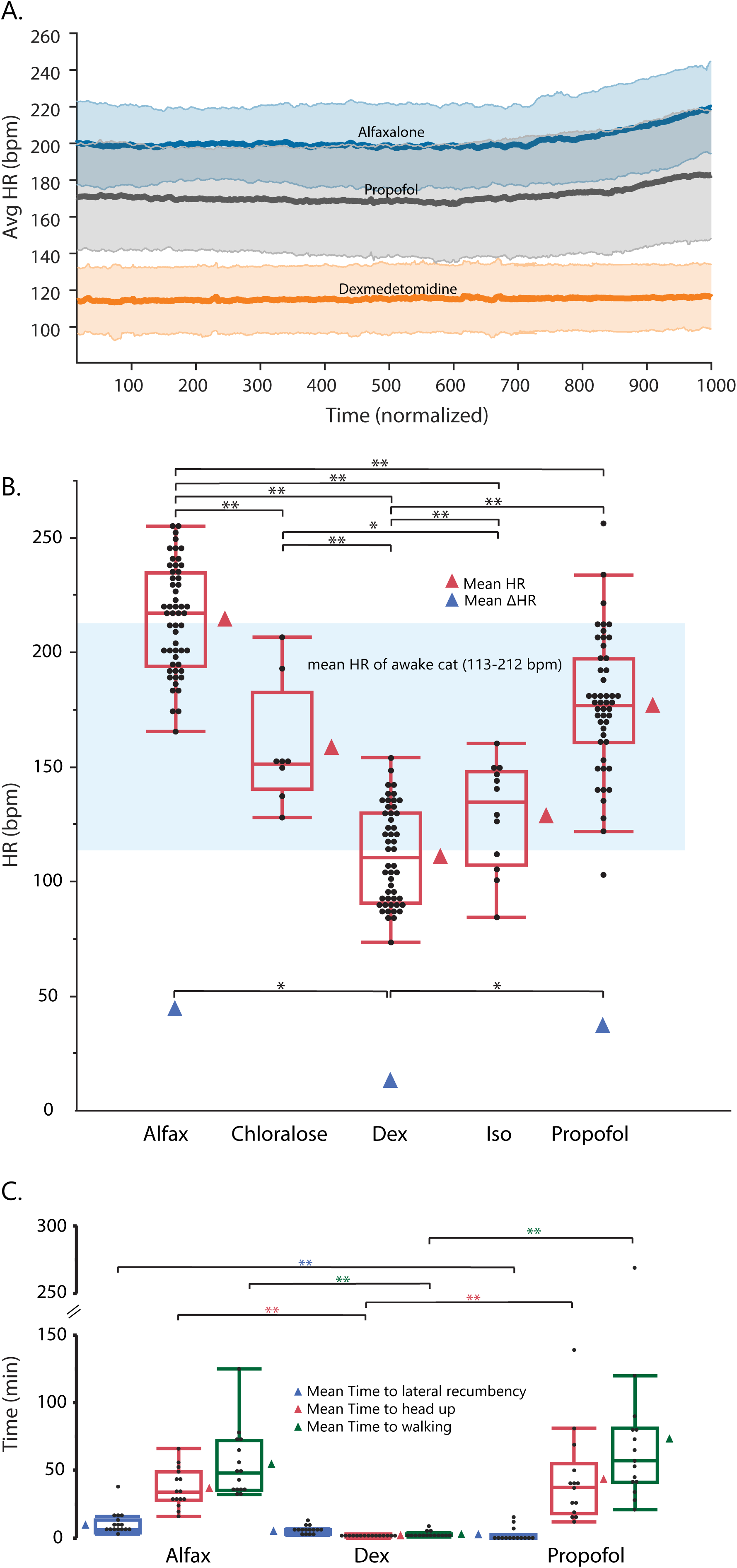
A. Averaged continuous HR of last 3 cats during CMG trials by agent, normalized by time. Solid lines represent means, shading around each mean represents standard deviations. B. Box plots of mean HR over all CMG trials by agent. ΔHR is shown for non-terminal agents only. Data represented as in Fig. 1. C. Box plots for time to induction (approximated by time to lateral recumbency), time to recovery (approximated by time to head up), and time to walking for non-terminal agents. All panels: significance: * p < 0.05, ** p < 0.01.

Time to lateral recumbency and time to head up and walking differed significantly between the non-terminal agents (Supplementary Table S5, Supplemental Table S6). The time to lateral recumbency was shortest under propofol due to IV administration (2.42 ± 1.62 minutes), and significantly shorter than alfaxalone (9.49 ± 1.49 minutes, p = 0.009). The time to head up and walking were shortest under dexmedetomidine (1.73 ± 6.07, 2.53 ± 10.58 minutes) due to use of the reversal agent atipamezole compared to alfaxalone (37.24 ± 6.10 [p < 0.001], 55.26 ± 10.64 [p < 0.001] minutes) and propofol (43.22 ± 6.28 [p < 0.001], 73.70 ± 10.64 [p < 0.001] minutes) (Fig. 3c). In general, reflexes showed greater anesthetic depth under dexmedetomidine compared to alfaxalone and propofol, although jaw tone was tighter under dexmedetomidine (Fig. 4).

**Figure 4:**
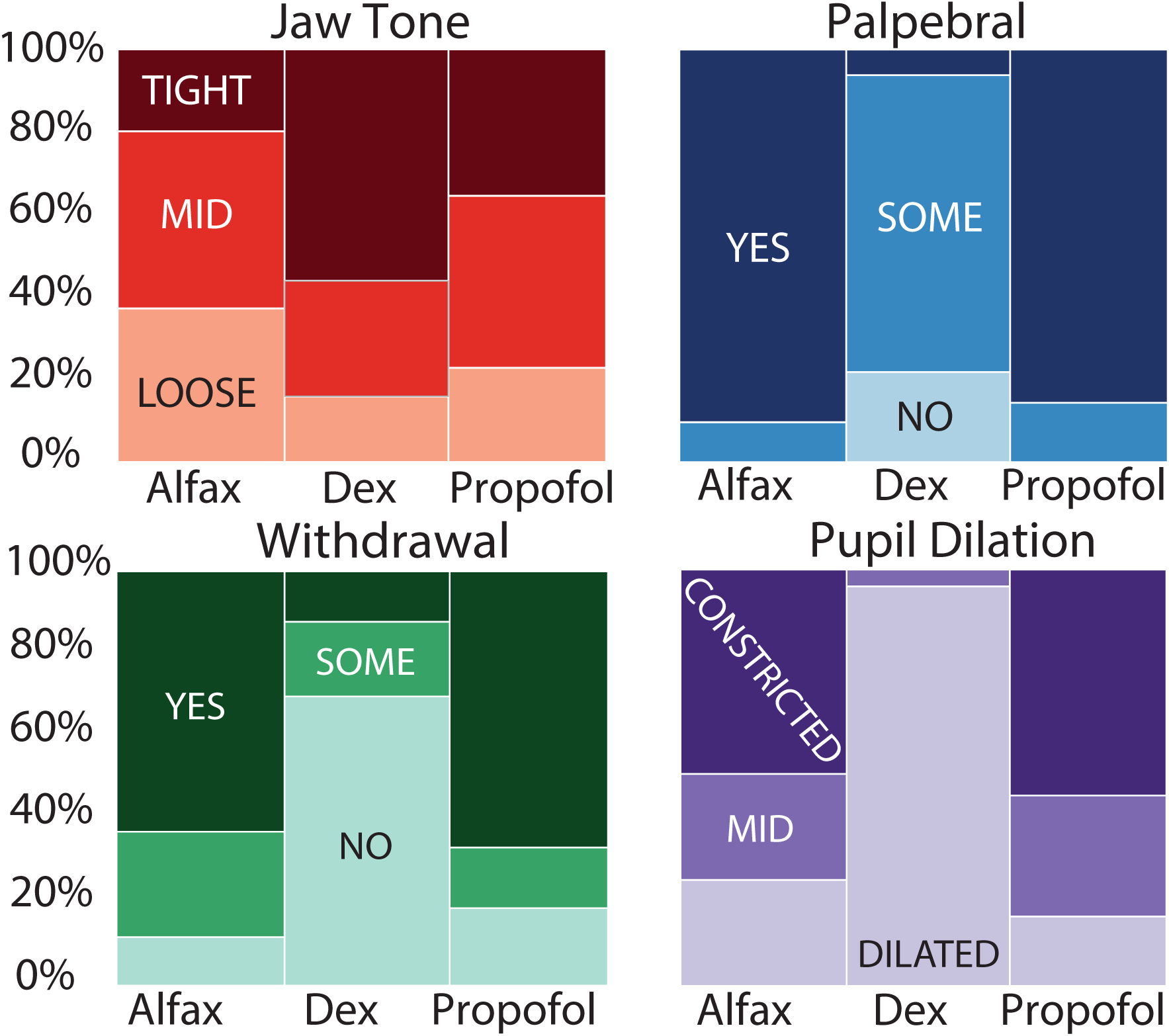
Reflex distribution by survival agents across all trials. Darker colors indicate a lighter plane of anesthesia.

### Adverse Consequences

Adverse consequences associated with VAP placement included surgical dehiscence (1/5), and port failure (1/5). In the cat with port failure, the last propofol trial was performed through isoflurane induction, in order to facilitate IV catheter placement. Dexmedetomidine caused vomiting immediately after injection in 7/15 (46.7%) of sessions. Cats did not vomit under other agents. Each cat vomited at least once under dexmedetomidine. One cat had an independent incidence of lethargy and vomiting unassociated with an anesthetic event, and was diagnosed with disseminated lymphoma at necropsy. One cat displayed significant bradycardia with 2^nd^ degree atrioventricular (AV) block arrhythmia under dexmedetomidine in 2 separate anesthetized events. Echocardiogram and subsequent necropsy of this cat suggested evidence of left ventricular diastolic dysfunction and myofibril degeneration and atrophy. Finally, 1/13 (7.7%) alfaxalone trials resulted in unexpected death. During recovery, after the CRI had been discontinued, one cat became apneic and was not able to be resuscitated.

## Discussion

In this study we demonstrated differences in urodynamic parameters and anesthetic depth at minimal doses needed to perform CMG testing in male cats. To our knowledge, this was the first study to examine CMG parameters under alfaxalone, propofol in isolation without another agent, or dexmedetomidine in comparison to the other agents used here. Our study incorporates a novel method of CMG assessment (bladder pressure slopes) in conjunction with traditional CMG parameters and careful evaluation of anesthetic depth. Together, our results provide recommendations for anesthetic choice based on study parameters of interest and logistic feasibility for CMG studies in male cats. Due to cross-species commonalities in response to anesthetic agents, select information from this study may be able to be guide preliminary anesthetic choice to other species, including humans. However, feasibility and effect on urodynamic and anesthetic parameters should be confirmed for the species in question.

Bladder capacity increases due to inhibition of central nervous system reflexes under anesthesia (post-operative urinary retention)^9^. In this study, the bladder increases seen under terminal agents (isoflurane and α-chloralose) (Fig. 1a) were strongly correlated with prolonged anesthesia (isoflurane [2-6 hours before CMG trials, 8-10 hours total], followed by α–chloralose [12-22 hours before CMG trials]) and repeated bladder fills for other experimental objectives. Although we did not see significant differences between survival agents, dexmedetomidine has been reported to cause polyuria through blockage of arginine-vasopressin release ^22,23^.

Maximum detrusor pressure [Pdet(max)] can also be used to assess bladder function ^15^. We reported Δpressure to account for any positioning differences between trials that may have affected baseline pressures. Consistent with previous studies examining Pdet(max) ^3,14^, we found that Δpressure was higher with propofol, and lower with isoflurane (Fig. 1b). While this may be attributed to the lighter plane of anesthesia under propofol (general anesthetics depresses Pdet(max) in a dose-dependent manner ^3,14^), a human study showed that even under comparable anesthetic depth, propofol showed less suppression of cortical somatosensory evoked potentials compared to isoflurane ^24^. Alternatively, the increase in Δpressure may be related to loss of function and initiation of the voiding reflex under propofol, as evidenced by bladder slopes drastically different from the awake state. While the results of propofol are difficult to interpret, isoflurane should be avoided in studies where Pdet(max) or Δpressure is a key parameter of interest.

Non-voiding contractions are phasic pressure increases seen during bladder filling ^25^ that may be linked to sensations relating to bladder volume ^26^. They are more likely and more frequently occur with urinary bladder pathologies ^25,27,28^. Consistent with our understanding that α-chloralose maintains spinal reflexes relative to other anesthetic agents ^2^, the greatest number of NVCs were observed with α-chloralose (Fig. 1c). Alfaxalone trials had the next greatest number of NVCs, and the most out of the non-terminal agents studied. Notably, dexmedetomidine almost completely eliminated NVCs compared to other agents. The α2 mediated analgesic properties of dexmedetomidine may block the sensation of bladder filling at the spinal or supra-spinal level ^16,29^. Clinically, dexmedetomidine has been used for analgesia and prevention of post-operative catheter-related bladder discomfort ^30^. However, awake CMG trials have significantly less NVCs than anesthetized trials ^12^, so absence of NVCs is not necessarily pathological. When NVCs are desired (ex: overactive bladder models), α-chloralose (non-survival) and alfaxalone (survival) can be considered, and dexmedetomidine should be avoided.

In this study CMG bladder pressure slopes differed between agents (Fig. 2), interpreted as previously described variable periods of passive filling and active contraction ^31^. Pressure slopes under dexmedetomidine (characterized by a flat filling period, followed by a sharp rise to the void event) are most similar to awake animal cystometrograms ^32,33^. In contrast, pressure slopes under propofol had the most obvious change from the awake slope, characterized by an early rapid rise and less obvious void event. Although there are no standardized criteria for defining passive filling and active contraction, we found propofol consistently reached a higher normalized pressure compared to dexmedetomidine across normalized time, with the most obvious separation noted between 900 - 950 normalized time, where the mean normalized pressure ± standard deviation range of dexmedetomidine is distinctly separate from the corresponding range for propofol (Fig. 2a). A previous rat study using a single anesthetic agent (tiletamine-zolazepam) at varying doses found lighter anesthesia was associated with more awake-like bladder curves ^32^. However, our study found dexmedetomidine (deeper plane of anesthesia) was associated with more awake-like bladder responses compared to GABA-ergic agonists with a lighter plane of anesthesia. This discrepancy suggests that suppression of bladder reflexes may not be directly correlated with others of anesthetic depth (anesthetic reflexes measured in this study, heart rate, ΔHR). The exact significance of bladder pressure slopes is unclear, but we recommend further investigation of bladder pressure response trends in future studies in association to anesthetic depth, pharmacokinetics, and preservation of spinal reflexes, as well as potential bladder pathology.

In addition to affecting CMG and urodynamic parameters, each agent offers unique logistical benefits and constraints. Dexmedetomidine (α_2_ agonist), conveniently administered through IM boluses (CRI only recommended for analgesia, not for sedation or anesthesia ^16^) and rapidly reversed through atipamazole, provides a convenient method of anesthetizing cats. Due to the nature of IM bolus administration, depth of anesthesia and therefore bladder function can vary as plasma levels of dexmedetomidine increase or decreases. Common adverse effects of dexmedetomidine include bradycardia, arrhythmias (AV block), and vomiting ^16^, all of which were noticed in our experiments.

Propofol and alfaxalone (GABA_A_ agonistics) are used clinically to both induce (IV) and maintain (CRI or repeat IV bolus) anesthesia ^11^. Maintenance of anesthesia using a CRI allows for steady-state drug plasma levels and a more stable plane of anesthesia. While propofol is limited to IV administration, alfaxalone can be given IM for single-agent anesthesia. Common side effects of both propofol and alfaxalone are apnea, although one study showed alfaxalone had less respiratory depression compared to propofol ^34^. Unexpectedly, we experienced a death during the recovery phase (most common time of anesthetic-related death ^35^) of one alfaxalone trial. To our knowledge, this was an isolated incident and not representative of the general safety of alfaxalone.

In this study, we were able to decease concentrations of GABA-ergic agonists below recommended labelled dosing in part due to the stable plane of anesthesia associated with CRI administration. This could be one explanation for the lighter plane of anesthesia (more preserved reflexes, higher average heart rate, and greater change in heart rate over trial) observed in GABA-ergic agonistics compared to dexmedetomidine (Figs. 3 and 4). However, another concurrent explanation for increased heart rate under the GABA-ergic agonists is hypotension with reflex tachycardia ^36^. This is in contrast with dexmedetomidine, which causes vasoconstriction, leading to hypertension and reflex bradycardia ^16^. Consistent with previous studies in dogs showing alfaxalone causes less cardiovascular depression than propofol ^36^, we observed the mean heart rate under alfaxalone (at or slightly above the upper limit of awake reference range ^37^) was significantly higher than under propofol (Fig. 3).

Isoflurane (exact mechanism of action unknown ^16,38^, but suspected to affect GABA receptors ^39^) is the primary inhalant anesthetic used in veterinary medicine, especially in prolonged invasive procedures ^16^.

Although in this study it was only used during terminal procedures, in clinical veterinary medicine isoflurane is commonly used during recovery procedures. Its inhaled route of administration allows dosing to be easily titratable to adjust the depth of anesthesia. However, inhaled agents are logistically more challenging, either requiring technical expertise to intubate the animal, or risking personnel exposure. Since recovery from isoflurane is rapid, and isoflurane has been previously used to facilitate propofol administration for urodynamic procedures ^40^, the effects of isoflurane on subsequent α-chloralose and propofol trials in this study are assumed to be minimal.

The non-survival hypnotic α-chloralose (mechanism unknown) provides light long-lasting anesthesia with minimal analgesia, and minimal effect on normal spinal and sympathetic reflex function ^2,11^. Unfortunately, α-chloralose is not recommended for survival studies due to involuntary excitement and prolonged recovery ^11^ although several studies have successfully recovered animals following α-chloralose anesthesia ^41,42^.

Limitations of this study include the following: 1) lack of direct comparison to awake CMG trials, 2) inability for full randomization due to logistical constraints (drug availability, ability to secure IV access, etc), although effort was made to spread out agents, so that each agent was tested once within the first 3 trials, once within the next 3 trials, and once within the final 3 trials, 3) inability to extrapolate data to female cats, 4) limited data for terminal agents (isoflurane and α-chloralose) due to nature of the experiments and the unexpected death of one animal, 5) lack of blinding in data analysis and experiments, especially for qualitative measures such as anesthetic reflexes, 6) lack of monitoring data other than heart rate due to measurement artifact and animal movement during trials, and 7) inability to control for different doses and routes of administration of a single drug due to logistical considerations and sample size. Future directions to address these limitations include additional routes and doses of single agents, concurrent CMG in awake cats for direct comparison to anesthetized cats, and filling of *ex-vivo* or post-mortem bladders to provide a truly passive bladder control.

## Methods

### Animals

Five adult, intact male domestic shorthair cats (7-18 months, 4-6 kg, Marshall BioResources, Inc., North Rose, NY) were included in this study. These animals were available from other study, and the cohort size is similar to other relevant studies ^3,14,17^. Animals were group housed in a temperature (72 ± 2 °F) and humidity (30 – 70%) controlled room on a 12:12 hour light/dark cycle with ad libitum access to water and fed a complete diet for adult cats (Purina One Urinary Tract Health Formula and Purina ProPlan EN, Nestle Purina, Neenah, WI). Cats were free of feline herpesvirus, calicivirus, panleukopenia, coronavirus, feline immunodeficiency virus, chlamydia, and toxoplasmosis. All experimental procedures were performed at an AAALAC accredited institution, according to the standards established by the Guide for the Care and Use of Laboratory Animals, and were approved by the University of Michigan Institutional Animal Care and Use Committee.

### Vascular Access Port

Each cat was implanted with a VAP (ClearPort Mid VAPs 5Fr with a rounded tip, 24” catheter; Access Technologies, Skokie, IL) to guarantee immediate IV access. IV access was essential for avoiding a second agent in propofol experiments, and useful in providing a constant rate infusion (CRI) in the alfaxalone and propofol experiments to maintain anesthesia. Cats were induced with dexmedetomidine (0.02 mg/kg, Dexmedseed, Dechra, Overland Park, KS), ketamine (6.6 mg/kg, Zetamine, Vet One, Boise, ID), and butorphanol (0.66 mg/kg, Torbugesic, Zoetis, Kalamazoo, MI), and maintained on isoflurane (0-2.5 %, Fluriso, Vet One, Boise, ID). Incisions were made on the left ventral neck and dorsal neck or back. A catheter was inserted and secured in the left jugular vein, and tunneled to a subcutaneous port secured to the muscle between the shoulder blades ^18,19^. The port was accessed with a non-coring Huber needle (Access Technologies, Skokie, IL), and flushed regularly (weekly, or after each use) with a locking solution (TCS Lock Solution, Access Technologies, Skokie, IL) or heparinized saline. Post-operative medications (buprenorphine [0.01-0.02 mg/kg, Par Pharmaceutical, Chestnut Ridge, NY], robenacoxib [6 mg/kg, Onsior, Elanco, Greenfield, IN] and famotidine [0.5 mg/kg, West-ward, Eatontown, NJ]) were given for 24 hours post-operatively. The cats were monitored daily until suture removal 7-14 days later.

### Anesthesia

Each cat was anesthetized at least three times (“sessions”) with each of three agents: (1) dexmedetomidine (0.02 – 0.04 mg/kg IM bolus; reversed with matched volume of atipamezole [Antisedan, Orion Corporation, Espoo, Finland] after session), (2) alfaxalone (5 mg/kg IM bolus + 0.08 mg/kg/min IV CRI, Alfaxan Multidose, Jurox Inc. Kansas City, MO), and (3) propofol (2 mg/kg IV bolus + 0.15 – 0.20 mg/kg/min IV CRI, Hospira, Inc., Lake Forest, IL). Effort was made to randomize sessions (each agent was tested once within the first 3 trials, once within the next 3 trials, and once within the final 3 trials), but full randomization could not be achieved due to logistical constraints (agent availability, repeat trials, etc.). Each session was on a different day, at least 2 days after the previous anesthetized session.

During terminal procedures in conjunction with other studies, cats were anesthetized with the same dexmedetomidine/ketamine/butorphanol combination as for the VAP surgery, and transitioned to isoflurane, then α-chloralose (70 mg/kg induction, 20 mg/kg maintenance every 4-6 hours or as needed [MilliporeSigma, Burlington, MA]), supplemented with 0.01 mg/kg buprenorphine ^20^.

### Cystometrograms

At least two CMG trials were conducted in each session. Warmed saline (41 °C) was pumped at 2 mL/min (AS50 infusion pump, Baxter, Deerfield, IL; or PHD 2000, Harvard Apparatus, Holliston, MA) through a fluid warmer (Hotline Fluid Warmer, Smiths Medical, Minneapolis, MN) and one lumen of a dual-lumen urethra catheter (Umbili-Cath™ 3.5 Fr Polyurethane UVC Catheter, Utah Medical, Midvale, UT) into the urinary bladder. For non-terminal sessions, sterile preparation of the perineal region and catheter was performed to mitigate against infection. The other lumen of the catheter was connected to a pressure transducer (V6402 pressure transducer, Smiths Medical, Minneapolis, MN) and Labchart (ADInstruments, Colorado Springs, CO) for recording. At bladder capacity a void event occurred and urine would leak around the catheter into a collecting tray (Fig. 5). The bladder was emptied and a break of at least 10 minutes was given between trials to allow the bladder to relax. Bladder capacity was calculated as the maximum of the total volume infused or the sum of urine collected from bladder and collecting tray. The variables analyzed for each CMG were bladder capacity, change in pressure from baseline to maximum (Δpressure), compliance (capacity/Δpressure), number and amplitude of non-voiding contractions (NVCs), and slopes of the bladder pressure (“pressure slopes”) over three intervals as defined below.

**Figure 5:**
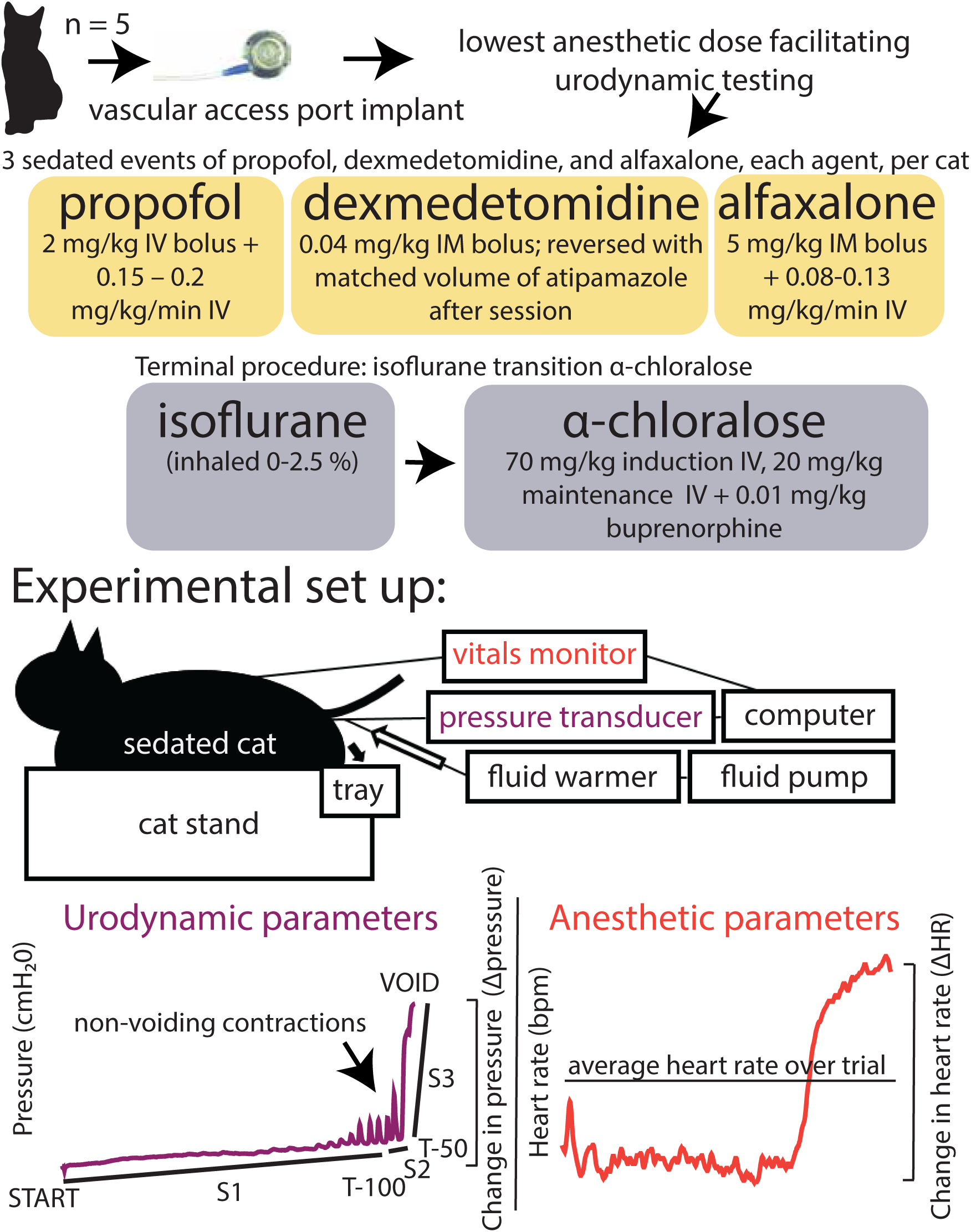
Experimental overview. Cats were implanted with VAPs and anesthetized at the lowest doses necessary to facilitate urodynamic testing. At least 3 sessions of propofol, dexmedetomidine, and alfaxalone each were conducted, in addition to a terminal experiment under isoflurane and α-chloralose in conjunction with another study. During anesthetized sessions at least 2 CMG trials were performed in each session to assess urodynamic parameters. Heart rate and reflexes were used to assess anesthetic depth, and time to induction/recovery was noted.

### Anesthetic Parameters

The time to lateral recumbency after initial agent injection and times to head up and walking (ten consecutive steps without falling) were recorded for each anesthetic session with non-terminal agents. During each session, vitals (heart rate [electrocardiogram or pulse oximetry], O_2_ hemoglobin saturation (pulse oximetry; tongue, lip, or ear), non-invasive blood pressure [forelimb] and temperature [rectal]) were monitored once every 15 minutes using a vitals monitor (Surgivet, Smiths Medical, Minneapolis, MN). Respiratory rate was taken manually. The average heart rate (mean HR) for a trial appeared to be a consistent indicator of anesthetic plane and was later included in statistical analysis. Other vitals either did not fluctuate across sessions or were subject to artifact and were not analyzed across sessions. During procedures for three of the five cats, a data logger was used for continuous vitals recording once per second and the calculation of change in heart rate over trial (ΔHR) in non-terminal studies. Reflexes (palpebral, withdrawal, jaw tone, pupil dilation) were checked before each CMG trial. Palpebral and withdrawal reflexes were categorized as “yes (strong withdrawal)”, “some (weak withdrawal)” or “no (no withdrawal)”. Jaw tone was categorized as “tight,” “mid” (intermediate), or “loose.” Pupil dilation was categorized as “constricted,” “mid” (intermediate), or “dilated.” In general, as anesthetic depth increases, palpebral and withdrawal reflexes decrease, jaw tone loosens, and pupils become more dilated ^21^.

### Data Acquisition and Analysis

Bladder pressure data was processed using MATLAB (R2019a, MathWorks, Natick, MA) and Excel (Microsoft, Redmond, WA). Bladder pressure was smoothed using a 1 second moving average. NVCs were identified in MATLAB using the findpeaks function, defining a peak as at least 3 cm H_2_O, at least 0.5 seconds away from another peak, and then manually confirmed. To account for the range of trial lengths, NVC counts were divided by the total length of the trial to calculate NVCs per second. To quantify overall bladder pressure trends seen between agents, pressure slopes were calculated for three distinct time periods: (1) infusion start to 100 seconds before void, (2) 100 to 50 seconds before void, and (3) last 50 seconds before void. To visualize the changes seen during slope calculations over different pressure and time ranges across cats and test sessions, the bladder pressure was normalized from 0 (start) to peak/void (1000) on the x (time) axis. After noting that pressures did not significantly fluctuate from resting baseline pressure until about 675 normalized time units, the 675-1000 normalized time unit range was targeted, and the pressure was normalized from 0 to 1 within that window. Continuous heart rate readings during non-terminal trials for the last 3 cats were also processed similarly, but over the entire length of the trial. Statistics were analyzed using JMP Pro 14.2.0 (SAS Institution Inc. Cary, NC), using a mixed model approach, setting anesthetic agent as a fixed variable and animal as a random variable. For analysis of bladder capacity, the cat weight was set as an additional fixed variable. Nominal categorical data was converted to ordinal categorical data in order to fit the mixed model. Yes/some/no, tight/mid/loose, and constricted/mid/dilated were converted to 0/0.5/1 respectively. Pairwise comparisons were made using Tukey’s HSD. Significance was defined as *p* < 0.05. Where relevant, data is reported as mean ± standard deviation.

## Data Availability

The datasets generated and analyzed during the current study and the MATLAB code used for data analysis are available in the Open Science Framework repository, (https://osf.io/8zjkp/ [DOI 10.17605/OSF.IO/8ZJKP]).

## Acknowledgements

We thank Zachariah Sperry, Ahmad Jiman, Elizabeth Bottorff, Hannah Parrish, and other members of the Peripheral Neural Engineering and Urodynamics (pNEURO) Lab for their assistance with surgeries, data collection, and analysis. We also thank Anna Skorupski, Samantha Eckley, Robert Sigler, and other members of the University of Michigan Unit for Laboratory Animal Medicine (ULAM) for their assistance with animal diagnostics and care. We thank the University of Michigan Consulting for Statistics, Computing and Analytics Research (CSCAR) for their advice on statistical analysis. This study was funded in part by the University of Michigan Cohen Comparative Medicine Research Award. No other funding sources contributed to this study.

## Author contributions

JJX, TMB, PAL and TM conceived and designed research. JJX, ZO, and EK developed logistics for performing experiment. JJX, ZY and EK performed experiments. JJX and ZY analyzed data. JJX, ZY, ZO and TMB interpreted results of experiments. JJX and ZY prepared figures. JJX, ZY, and TMB drafted manuscript. JJX, ZY, ZO, PAL, TM and TMB edited and revised manuscript. All authors approved the final version of the manuscript.

