## Supplemental Tables for "Anesthetic agents affect urodynamic parameters and anesthetic depth at doses necessary to facilitate preclinical testing in felines"

Table S1: Table of cats and analyzed trials. Data reported as: total trials (# of sessions).

| Cat/Agent | Alfaxalone | Propofol | Dexmedetomidine | Isoflurane | α-chloralose |
|---|---|---|---|---|---|
| Cat #1 | 8 (3) | 9 (3) | 11 (3) | 3 (1) | 2 (1) |
| Cat #2 | 11 (4) | 11 (4) | 9 (3) | 3 (1) | 3 (1) |
| Cat #3 | 10 (3) | 10 (3) | 9 (3) | N/A | N/A |
| Cat #4 | 10 (3) | 9 (3) | 9 (3) | 3 (1) | 3 (1) |
| Cat #5 | 12 (3) | 9 (3) | 10 (3) | 3 (1) | 3 (1) |

Table S2: Least squares means estimates and 95% confidence intervals for CMG parameters.

| | Alfaxalone | α-chloralose | Dexmedetomidine | Isoflurane | Propofol |
|---|---|---|---|---|---|
| BLADDER CAPACITY (ml) | | | | | |
| Estimate | 46.72 | 59.61 | 42.93 | 54.54 | 44.18 |
| Std Error | 8.56 | 9.18 | 8.56 | 9.11 | 8.56 |
| Lower 95% | 12.11 | 29.49 | 8.38 | 24.10 | 9.71 |
| Upper 95% | 81.33 | 89.74 | 77.48 | 84.97 | 78.65 |
| ΔPRESSURE (cmH20) | | | | | |
| Estimate | 97.98 | 108.08 | 101.53 | 84.24 | 116.53 |
| Std Error | 9.53 | 13.51 | 9.61 | 13.15 | 9.61 |
| Lower 95% | 74.78 | 80.15 | 78.33 | 56.91 | 93.32 |
| Upper 95% | 121.17 | 136.01 | 124.74 | 111.57 | 139.74 |
| COMPLIANCE (ml/cm$H_2$0) | | | | | |
| Estimate | 0.64 | 0.58 | 0.48 | 0.61 | 0.49 |
| Std Error | 0.12 | 0.16 | 0.12 | 0.15 | 0.12 |
| Lower 95% | 0.33 | 0.24 | 0.18 | 0.28 | 0.18 |
| Upper 95% | 0.95 | 0.91 | 0.79 | 0.94 | 0.80 |
| NVC (# of NVC/s) | | | | | |
| Estimate | 0.012 | 0.033 | 0.002 | 0.008 | 0.010 |
| Std Error | 0.003 | 0.005 | 0.003 | 0.005 | 0.003 |
| Lower 95% | 0.006 | 0.023 | -0.004 | -0.002 | 0.003 |
| Upper 95% | 0.019 | 0.043 | 0.009 | 0.017 | 0.016 |
| NVC AMPLITUDE (cm$H_2$0) | | | | | |
| Estimate | 6.93 | 7.93 | 5.42 | 7.05 | 4.94 |
| Std Error | 0.99 | 1.41 | 1.03 | 1.48 | 0.99 |
| Lower 95% | 4.56 | 5.02 | 3.03 | 4.03 | 2.57 |
| Upper 95% | 9.30 | 10.84 | 7.81 | 10.07 | 7.31 |
| SLOPE 1: start to T-100 (cm$H_2$0/s) | | | | | |
| Estimate | 0.03 | 0.03 | 0.02 | 0.03 | 0.06 |

| | | | | | |
|---|---|---|---|---|---|
| Std Error | 0.01 | 0.01 | 0.01 | 0.01 | 0.01 |
| Lower 95% | 0.00 | 0.00 | 0.00 | 0.01 | 0.04 |
| Upper 95% | 0.05 | 0.06 | 0.05 | 0.06 | 0.09 |
| SLOPE 2: T-100 to T-50 ($cmH_20/s$) | | | | | |
| Estimate | 0.27 | 0.51 | 0.17 | 0.28 | 0.36 |
| Std Error | 0.05 | 0.08 | 0.05 | 0.08 | 0.05 |
| Lower 95% | 0.16 | 0.34 | 0.06 | 0.12 | 0.24 |
| Upper 95% | 0.39 | 0.67 | 0.29 | 0.43 | 0.47 |
| SLOPE 3: T-50 to void ($cmH_20/s$) | | | | | |
| Estimate | 1.10 | 0.63 | 1.45 | 0.57 | 0.65 |
| Std Error | 0.11 | 0.18 | 0.11 | 0.17 | 0.11 |
| Lower 95% | 0.85 | 0.27 | 1.19 | 0.23 | 0.40 |
| Upper 95% | 1.36 | 0.98 | 1.70 | 0.92 | 0.91 |

Table S3: p values of Tukey's HSD all pairwise comparisons for each CMG parameter. Significance: * p < 0.05, ** p < 0.01.

| Agent 1 – Agent 2 | | p values | | | | | | | |
|---|---|---|---|---|---|---|---|---|---|
| Agent 1 | Agent 2 | Bladder capacity | ΔPressure | Compliance | NVC (#/s) | NVC amplitude | Slope 1 | Slope 2 | Slope 3 |
| Alfax | Chloralose | 0.0182* | 0.9138 | 0.9868 | 0.0003** | 0.9356 | 0.9987 | 0.0295* | 0.0449* |
| Alfax | Dex | 0.5598 | 0.9875 | 0.2117 | 0.0084** | 0.3635 | 0.9999 | 0.2091 | 0.0094** |
| Alfax | Iso | 0.2943 | 0.7521 | 0.9995 | 0.8543 | 1.0000 | 0.9522 | 1.0000 | 0.0129* |
| Alfax | Propofol | 0.8520 | 0.0749 | 0.2538 | 0.8941 | 0.0881 | <.0001** | 0.4156 | 0.0002** |
| Chloralose | Dex | 0.0008** | 0.9820 | 0.9379 | <.0001** | 0.3123 | 0.9967 | 0.0004** | <.0001** |
| Chloralose | Iso | 0.8612 | 0.4920 | 0.9993 | 0.0004** | 0.9847 | 0.9973 | 0.1376 | 0.9990 |
| Chloralose | Propofol | 0.0026** | 0.9547 | 0.9531 | <.0001** | 0.1411 | 0.0724 | 0.3240 | 0.9999 |
| Dex | Iso | 0.0338* | 0.5628 | 0.8055 | 0.8115 | 0.7596 | 0.9321 | 0.6539 | <.0001** |
| Dex | Propofol | 0.9873 | 0.2378 | 1.0000 | 0.1137 | 0.9773 | <.0001** | 0.0018** | <.0001** |
| Iso | Propofol | 0.0790 | 0.0449* | 0.8371 | 0.9925 | 0.5194 | 0.1567 | 0.8371 | 0.9886 |

Table S4: Least squares means estimates and 95% confidence intervals for anesthetic parameters.

| | Alfaxalone | α-chloralose | Dexmedetomidine | Isoflurane | Propofol |
|---|---|---|---|---|---|
| HEART RATE (beats per minute) | | | | | |
| Estimate | 215.16 | 163.29 | 110.66 | 130.73 | 176.72 |
| Std Error | 5.69 | 9.49 | 5.75 | 8.18 | 5.77 |
| Lower 95% | 201.38 | 144.13 | 96.87 | 113.89 | 162.92 |
| Upper 95% | 228.94 | 182.44 | 124.45 | 147.58 | 190.52 |
| ΔHR (beats per minute) | | | | | |
| Estimate | 45.36 | | 13.79 | | 37.50 |
| Std Error | 6.65 | | 6.80 | | 7.05 |
| Lower 95% | 27.42 | | -4.01 | | 19.81 |
| Upper 95% | 63.31 | | 31.59 | | 55.19 |
| Time to lateral recumbency (min) | | | | | |
| Estimate | 9.49 | | 5.00 | | 2.42 |
| Std Error | 1.49 | | 1.54 | | 1.62 |
| Lower 95% | 6.41 | | 1.85 | | -0.92 |
| Upper 95% | 12.56 | | 8.15 | | 5.76 |
| Time to head up (min) | | | | | |
| Estimate | 37.24 | | 1.73 | | 43.22 |
| Std Error | 6.10 | | 6.07 | | 6.28 |
| Lower 95% | 24.10 | | -11.33 | | 29.81 |
| Upper 95% | 50.38 | | 14.80 | | 56.64 |
| Time to walking (min) | | | | | |
| Estimate | 55.26 | | 2.53 | | 73.70 |
| Std Error | 10.64 | | 10.58 | | 10.64 |
| Lower 95% | 32.72 | | -19.86 | | 51.16 |
| Upper 95% | 77.80 | | 24.92 | | 96.24 |

Table S5: p values of Tukey's HSD all pairwise comparisons for each anesthetic parameter. Not all parameters were assessed for terminal agents (isoflurane and α-chloralose). Significance: * $p < 0.05$, ** $p < 0.01$.

| Agent 1 – Agent 2 | | p values | | | | |
|---|---|---|---|---|---|---|
| Agent 1 | Agent 2 | HR | Δ HR | Time to lateral recumbency | Time to head up | Time to walking |
| Alfax | Chloralose | <.0001* | | | | |
| Alfax | Dex | <.0001* | <.0001* | 0.1206 | <.0001* | 0.0009* |
| Alfax | Iso | <.0001* | | | | |
| Alfax | Propofol | <.0001* | 0.7766 | 0.0093* | 0.7039 | 0.3585 |
| Chloralose | Dex | <.0001* | | | | |
| Chloralose | Iso | 0.0187* | | | | |
| Chloralose | Propofol | 0.5539 | | | | |
| Dex | Iso | 0.0561 | | | | |
| Dex | Propofol | <.0001* | 0.0082* | 0.5086 | <.0001* | <.0001* |
| Iso | Propofol | <.0001* | | | | |

Table S6: p values of Tukey's HSD pairwise comparisons for reflexes during non-terminal trials. Significance: * p < 0.05, ** p < 0.01.

| Anesthetic agents | | p value | | | |
|---|---|---|---|---|---|
| Agent 1 | Agent 2 | Jaw Tone | Withdrawal | Palpebral | Pupil dilation |
| Alfaxalone | Dexmedetomidine | 0.0003* | <.0001* | <.0001* | <.0001* |
| Alfaxalone | Propofol | 0.0753 | 0.9604 | 0.8070 | 0.4890 |
| Dexmedetomidine | Propofol | 0.1965 | <.0001* | <.0001* | <.0001* |
